# Single-molecule dynamics and genome-wide transcriptomics reveal that NF-κB (p65)-DNA binding times can be decoupled from transcriptional activation

**DOI:** 10.1101/255380

**Authors:** Andrea Callegari, Christian Sieben, Alexander Benke, David M. Suter, Beat Fierz, Davide Mazza, Suliana Manley

## Abstract

Transcription factors (TFs) regulate gene expression in both prokaryotes and eukaryotes by recognizing and binding to specific DNA promoter sequences. In higher eukaryotes, it remains unclear how the duration of TF binding to DNA relates to downstream transcriptional output. Here, we address this question for the transcriptional activator NF-κB (p65), by live-cell single molecule imaging of TF-DNA binding kinetics and genome-wide quantification of p65-mediated transcription. We used mutants of p65, perturbing either the DNA binding domain (DBD) or the protein-protein transactivation domain (TAD). We found that p65-DNA binding time was predominantly determined by its DBD and directly correlated with its transcriptional output as long as the TAD is intact. Surprisingly, mutation or deletion of the TAD did not modify p65-DNA binding stability, suggesting that the p65 TAD generally contributes neither to the assembly of an “enhanceosome,” nor to the active removal of p65 from putative specific binding sites. However, TAD removal did reduce p65-mediated transcriptional activation, indicating that protein-protein interactions act to translate the long-lived p65-DNA binding into productive transcription.

**Author Summary:** To control transcription of a certain gene or a group of genes, both eukaryotes and prokaryotes express specialized proteins, transcription factors (TFs). During gene activation, TFs bind gene promotor sequences to recruit the transcriptional machinery including DNA polymerase II. TFs are often multi-subunit proteins containing a DNA-binding domain (DBD) as well as a protein-protein interaction interface. It was suggested that the duration of a TF-DNA binding event 1) depends on these two subunits and 2) dictates the outcome, i.e. the amount of mRNA produced from an activated gene. We set out to address these hypotheses using the transcriptional activator NF-κB (p65) as well as a number of mutants affecting different functional subunits. Using a combination of live-cell microscopy and RNA sequencing, we show that p65 DNA-binding time indeed correlates with the transcriptional output, but that this relationship depends on, and hence can be uncoupled by altering, the protein-protein interaction capacity. Our results suggest that, while p65 DNA binding times are dominated by the DBD, a transcriptional output can only be achieved with a functional protein-protein interaction subunit.

## Introduction

Transcription factors (TFs) are fundamental regulatory components of transcription in both prokaryotes and eukaryotes, which can activate or repress the expression of specific genes. The NF-κB family of TFs, universal among nearly all animal cell types, is involved in many signaling pathways and when dysregulated can contribute to several pathologies, including cancer and inflammatory diseases [1]. This is exemplified by the RELA (v-rel reticuloendotheliosis viral oncogene homolog A), or p65 TF, which is implicated in regulating the activation of ~150 genes involved in wide-ranging functions from immune response to metabolism [1]. In its most prevalent form, p65 forms a stable heterodimer with p50 in the cytoplasm [2] (Fig 1A). Upon stimulation, the activated heterodimer translocates into the nucleus [3]. The heterodimer interacts with target DNA regulatory elements through a conserved Rel homology region (RHR) [4–6]. Following DNA binding, p65-mediated transcriptional activation is controlled by two trans-activation domains (TADs), TAD1 and TAD2 [7]. Co-regulators of transcription are recruited at the promoters of target genes via protein-protein interactions mediated by TAD1 and TAD2, eventually leading to the recruitment of RNA polymerase II (RNA pol-II) and subsequent activation of gene expression [8]. Deletion of one or both TADs has been shown to heavily impair p65-dependent transcriptional activation, suggesting a dominant-negative effect of such truncation mutants [7].

**Fig 1.**
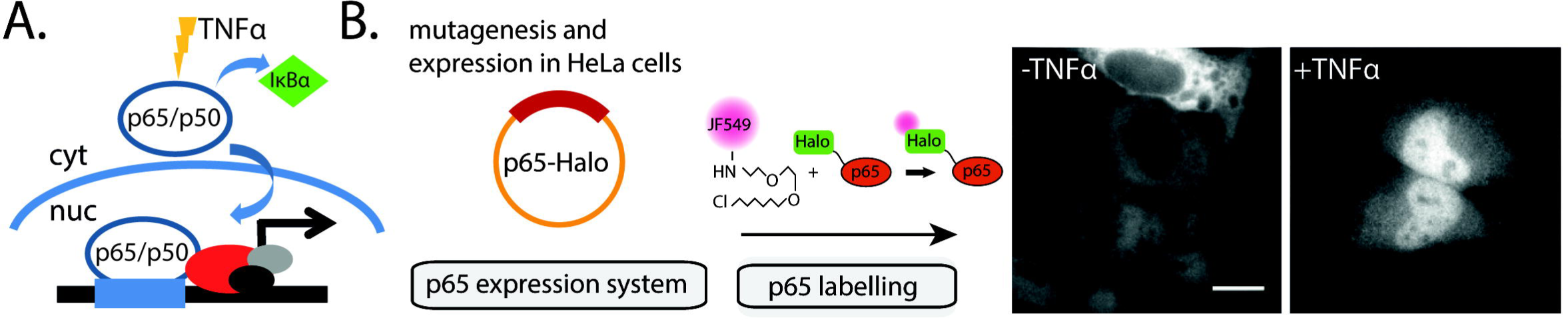
**NFκB-p65 mode of action and experimental setup. (A)** p65 forms a heterodimeric complex with p50 in the cytosol (cyt). The complex is bound by the cytosolic inhibitor *IκBa* that prevents its translocation into the nucleus. Following TNF-α stimulation and *IκBα* dissociation, p65/p50 translocate into the nucleus allowing subsequent DNA binding and gene activation. **(B)** We used a copy of the human p65 fused to a Halo tag as a basis for our mutational expression system. P65-Halo constructs were then expressed in HeLa cells and fluorescently labelled with a JF549 Halo ligand. Upon TNF-α stimulation, labelled p65-Halo translocates into the nucleus. Scale bar: 10 μm.

There is increasing evidence from biochemistry and live-cell single molecule imaging that in general the duration of binding events of TFs to responsive elements (RE) correlates with transcriptional activity [9–11]. However, as in the case of p65, it remains largely unknown whether protein-protein interactions mediated by p65 TADs can stabilize DNA binding, and thus lead to higher transcriptional activity. Such a stabilization would be expected according to the enhanceosome model, for which a stable higher-order complex formed by NF-κB and additional cofactors interacting with its TADs is assembled at the IFN-β1 regulatory element. The enhanceosome model was challenged by a subsequent study in which the duration of p65-DNA binding was measured at genetically engineered arrays in cells and found to be very transient [12], an observation incompatible with the formation of stable complexes. Instead, perhaps protein-protein interactions mediated by the p65 TADs could destabilize its binding to DNA, and TAD-mediated transactivation would rather be responsible for actively displacing p65 from chromatin. As this live-cell study was performed on artificial arrays of p65 binding sites, the role of TAD-mediated protein-protein interactions for p65-binding stability in the genomic context still remains unexplored.

Here, we combined single-molecule live-cell imaging and genome-wide transcriptomics of wild-type p65 (wt-p65), TAD truncation mutants and p65 DNA-binding affinity mutants to elucidate the role of these domains in the stability of p65 binding and on downstream transcriptional activity. We established point mutants to the DNA-binding domain to modulate p65-DNA binding affinity. We found that the lifetime of binding events of p65 mutants to chromatin in living cells correlated with their reported *in vitro* binding affinities and genome-wide transcriptional activity. We next examined the effects of TAD deletion mutants. We found that these mutants had DNA-binding kinetics comparable to wt-p65. However, whole transcriptome profiling revealed that TAD truncated forms of p65 did have impaired transactivation capability, suggesting that TAD-mediated protein-protein interactions serve the role of translating longer-lived p65-DNA binding into transcriptionally productive events.

## Results

We carried out a p65 DNA-binding kinetics and genome-wide transcriptomic study using a carboxy-terminal Halo-Tag [13] fusion construct of the human p65 (p65-Halo) (Fig 1b). To fluorescently label p65, HeLa cells were transiently transfected with p65-Halo and incubated with Halo-JF549 [14] (Fig 1b). In a large majority of transfected cells (~90%), the labeled p65-Halo was enriched in the cytosol and excluded from the nucleus (Fig 1b, panel “-TNFα”). After 30 minutes of stimulation with TNFα, p65-Halo translocated from the cytosol into the nucleus in ~73% of the cells (Fig 1b, panel “+TNFα”), showing that it was responsive to TNFα treatment.

We also tested the p65-Halo fusion protein for its ability to transactivate two well-known p65 target genes, *NFKBIA* and *Ccl2*, either in the presence or absence of TNFα stimulation (S1 Fig). Ectopically expressed p65-Halo upregulated the expression of both genes above their endogenous levels in non-stimulated cells (*p* < 0.05). Furthermore, upon TNFα stimulation, *NFKBIA* showed a significantly (*p* < 0.05), increased level of expression, whereas *Ccl2* upregulation was not significantly different (*p* > 0.05) from non-stimulated cells. This is likely due to two synergic factors, that is, the high expression levels of *Ccl2* in the presence of overexpressed p65 and the slower activation rate of *Ccl2* as compared to *NFKBIA* [15, 16].

We further verified the interaction of p65-Halo with its consensus DNA sequence (S2 Fig) by using an electrophoretic mobility shift assay (EMSA). JF549-labeled p65-Halo was purified (Materials and Methods) and incubated with Atto647N-labeled consensus oligonucleotide before electrophoretic separation under non-denaturing conditions (Materials and Methods). Increasing concentrations of p65-Halo enhanced the shifted fraction of labeled oligonucleotide, confirming the ability of the fusion protein to bind *in vitro* to its specific consensus sequence (Fig 1c, S2 Fig).

### The role of p65-DNA affinity in determining its nuclear DNA binding time

We performed 2D single-molecule tracking (SMT) of individual, JF549-labeled p65-Halo molecules in the nucleus of live HeLa cells after stimulation with TNFα. We excited the sample with a highly inclined and laminated laser illumination (HILO) to minimize background fluorescence from out-of-focus p65 molecules [17] (Materials and Methods). Further, using stroboscopic laser excitation (*t_int_* = 5 *ms*; *t_gap_* = 95 *ms*; power ~1 kW cm^−2^), we could minimize photobleaching, allowing us to record long (seconds) trajectories from both static and mobile p65 molecules (Fig 2a). To selectively identify p65 molecules bound to chromatin, we used the histone subunit H2B fused to Halo tag as an “immobile” control to define an upper threshold for the displacement _1-2_ between two consecutive frames. We found that ~99% of H2B displacements were below *r_max_* = 435 *nm* (Fig 2a and S3 Fig). Each p65 frame-to-frame displacement satisfying *r* < *r_max_* was further required to last at least 10 frames to minimize the probability that slowly diffusing molecules would affect the calculated *t_b_* [18]. The binding time *t_b_* of each DNA-bound p65 single molecule was then directly measured as the number of frames the fluorescence stayed “on” until disappearance. We found that *t_b_* of DNA-bound p65 molecules could not be described by a single-exponential decay model (S4 Fig). However, a bi-exponential decay model was in good agreement with our photobleaching-corrected data (Fig 2c and S4 Fig), with lifetimes of 0.53 s and 4.1 s for the short- and long-lived populations of p65-Halo wild-type molecules (wt-p65), respectively (Fig 2d). Short- and long-lived populations corresponded to ~95.7% and ~4.3% of wt-p65 DNA-bound molecules (Fig 2d).

**Fig 2.**
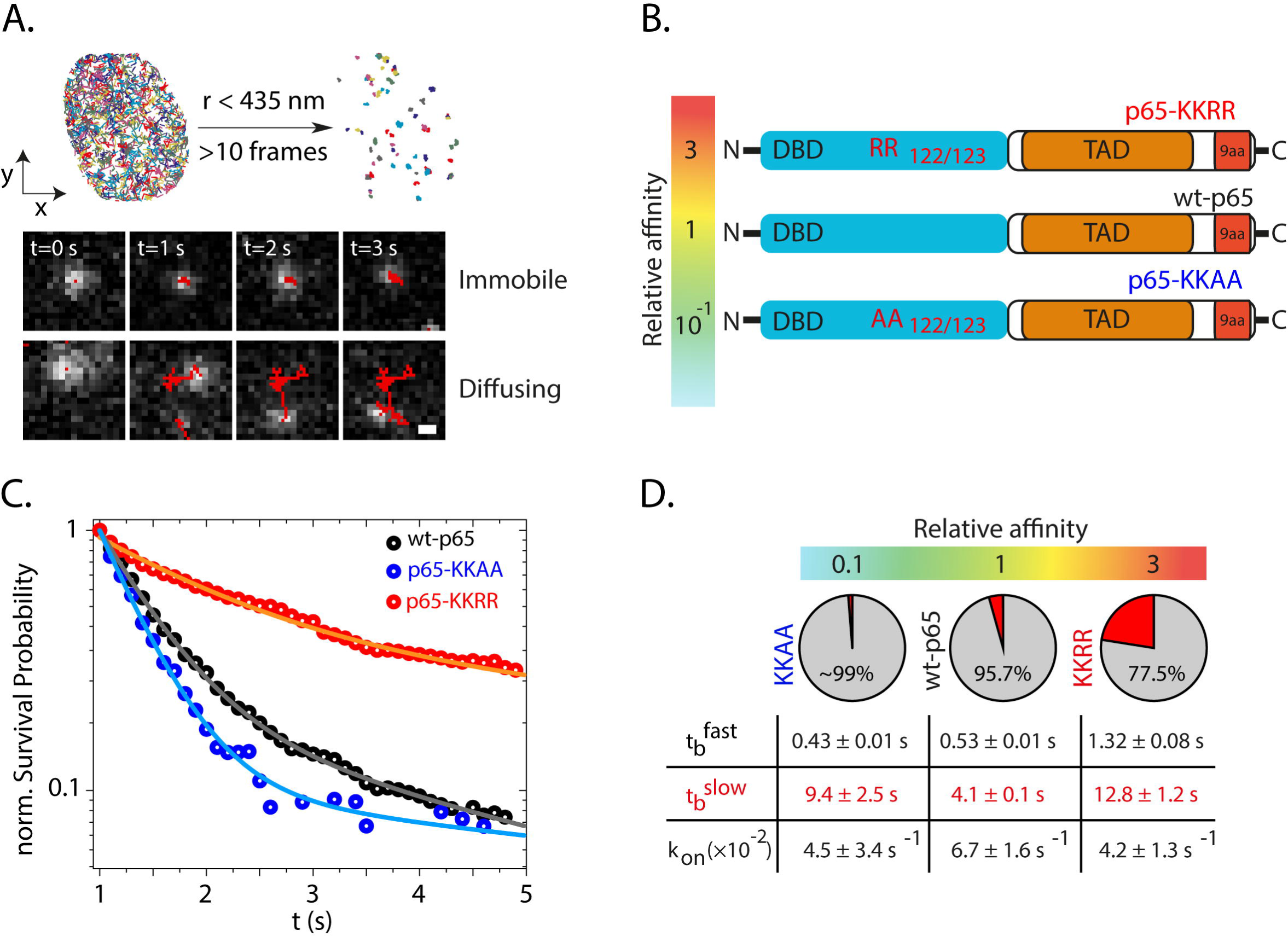
**DNA binding time of p65 DNA affinity mutants. (A)** Single particle tracking (SPT) was performed after TNF-α stimulation and p65-Halo translocation into the nucleus. All recorded trajectories were filtered based on a spatial threshold established using an immobile control (Histone subunit H2B, see Fig. S3). After filtering out mobile molecules and correction of photobleaching, the DNA binding time [35] could be estimated from the length of each individual trajectory. (**B**) Schematic overview of p65 affinity mutants. **(C)** Normalized survival probability plots (1-CDF plot) of the DNA-bound fraction for wt-p65 as well as the DNA affinity mutants KKAA and KKRR. The distributions were fitted using a bi-exponential function revealing the fast (t_b_fast) and slow (t_b_slow) DNA binding times. **(D)** Summary of the obtained fitting parameters together with the relative DNA dissociation constant K_D_ for each construct. The pie chart shows the fraction of events associated to the fast (grey) or slow [36] binding time. *K_on_\** was obtained from single step displacement histograms as described in Methods. While all constructs exhibit similar k_on_* as well as t_b_fast, the slow binding time t_b_slow correlates with the DNA affinity.

To modulate the affinity of p65 for DNA, we performed single-point mutagenesis within the p65 DNA-binding domain (DBD) [19]. We generated two DNA binding affinity mutants, p65-KKAA-Halo and p65-KKRR-Halo, corresponding to lower (relative *K_D_* = 0.1) or higher (relative *K_D_* = 3) *in vitro* binding affinities as compared to wt-p65 [19]. We expressed each mutant in HeLa cells and initially performed fluorescence recovery after photobleaching (FRAP, S5 Fig). As expected, the lower DNA-binding affinity mutant p65-KKAA showed faster recovery as compared to wt-p65 (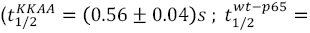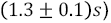) while p65-KKRR displayed comparable recovery dynamics (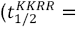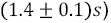) to wt-p65.

SMT followed by bi-exponential fitting of the survival probability distributions identified two distinct binding components for both affinity mutants as observed for wt-p65 (Fig 1c, d). Consistent with the estimated bound fractions from FRAP, p65-KKAA showed only a residual ~1% of long-lived binding events 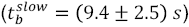, while the p65-KKRR variant displayed a significantly higher fraction of long-lived binding events (~22.5%) associated with longer binding times 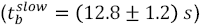. To provide quantitative estimates of the fraction of p65 molecules involved in binding, we repeated the SMT at faster frame rates (*t_int_* = 5 *ms*; *t_gap_* = 15 *ms*) and analyzed the resulting tracks by fitting the distribution of displacements between consecutive frames (Δ*t* = 20 *ms*) using a three-component diffusive model [20] (Equation 1). The fraction of p65 molecules corresponding to the slowest diffusing component matched the diffusivity coefficient of the histone subunit H2B (~0.04 μm^2^ s^−1^) and identified the BF of p65 molecules. We noted that p65-KKRR displayed a significantly higher BF (*BF^KKRR^*~30%) than p65-KKAA (*BF^KKAA^*~4%) and wt-p65 (*BF^wt-p65^*~21%; S5 Fig) which well explains the higher immobile fractions in FRAP recovery curves (S5 Fig).

### The transcriptional activation potential of p65 mutants

The transcriptional activation potential of p65-Halo mutants was estimated by measuring transcriptome-wide gene expression levels (RNAseq; see Methods). RNAseq analysis allowed us to identify differentially expressed genes by comparing stimulated (+TNFα) and non-stimulated cells (-TNFα). Using a false-discovery rate (FDR) lower than 0.1 (Materials and Methods), a total of 1080 genes were scored as differentially expressed (Fig 3a). Of these, we selected only genes directly bound by p65 on the basis of deposited ChIP-seq data (ENCODE database). Among the remaining 215 genes, we identified 15 well-characterized p65 targets, including *FAS, IL23A* and *TRAF1*. The relative fold-changes (FC) of expression of the 215 p65-target genes were then computed for each generated mutant by normalizing against the gene expression levels observed in non-transfected cells (NT) (Fig 3a). This analysis was complemented by determining the *z*-score of gene expression levels and visualized using a heat-map (Fig 3b). Results obtained with both approaches identified p65-KKAA as a loss-of-function and p65-KKRR as a gain-of-function mutant (Fig 3b-d).

**Fig 3.**
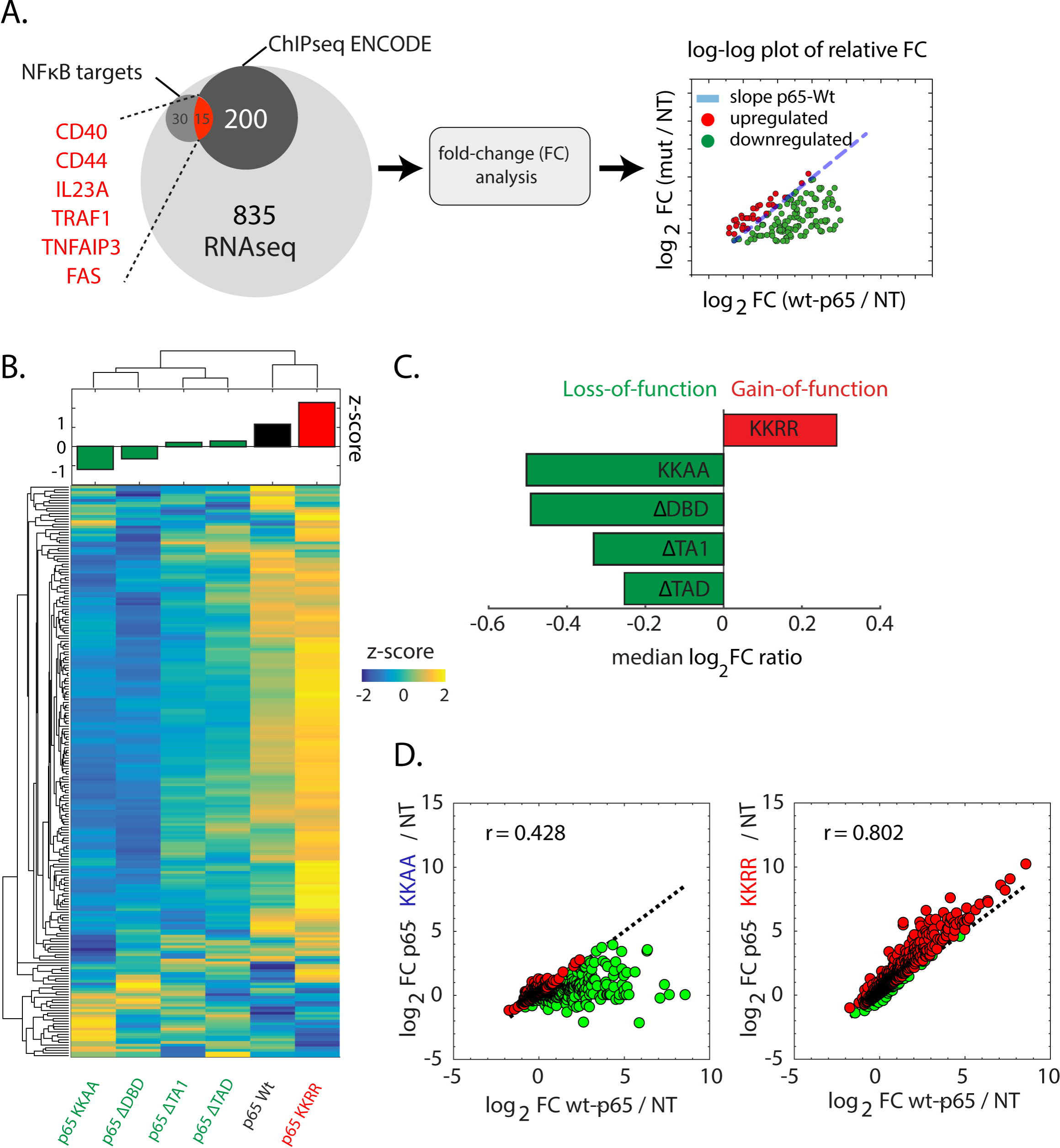
**Transcriptional activity of p65 DNA affinity mutants. (A)** We assessed the level of gene activation using RNA sequencing (RNA-Seq). To this end, total mRNA was isolated and sequenced using next-generation sequencing. The total initial set of 1080 genes was cross-referenced using the Chip-Seq ENCODE database, providing a subset of 215 direct interacting genes. A second subset of 45 genes consist of known *NFκB* regulated genes. For each p65 mutant, the fold-change (FC) expression above the non-transfected (NT) control was calculated and for each gene compared with wt-p65 using a log-log FC plot. As a general discrimination between up- and downregulated genes compared to wt-p65, we calculated the logFC ratio. Values with logFC ratio>1 are marked upregulated (red), those with logFC ratio<1 are marked downregulated (green). **(B)** FC values were standardized (per gene) and the different conditions clustered hierarchically to identify similarities. Interestingly, wt-p65 co-clusters with p65-KKRR, which also shows the highest average z-score (**B**, top plot) and was identified as the only gain-of-function mutant **(C)**. The two transactivation mutants as well as the low affinity mutant and p65-ΔDNA also co-cluster highlighting their functional similarity. **(C)** Classification of each p65 variant based on the logFC ratio estimator, showing that p65-KKRR (i.e. with higher DNA affinity) represents the only gain-of-function mutant. **(D)** RNA-Seq analysis comparing transcriptional activation of p65-KKAA and p65-KKRR with wt-p65. p65-KKAA shows very weak correlation with wt-p65 as well as a strongly reduced gene activation (logFC ratio = −0.47) indicating a loss of gene specificity as well as activation potential. In contrast, p65-KKRR shows higher correlation as well as an increased gene activation (logFC ratio = 0.28).

### DNA-binding time and transcriptional activation potential of p65 truncation mutants

To investigate the role of protein-protein interactions on the p65 DNA binding time and downstream transcriptional activation, we generated two additional truncation mutants, lacking one (p65-DTAD1) or both TADs (p65-DTAD; Fig 4a). We performed SMT and RNAseq on these TAD mutants to retrieve binding kinetics (Fig 4b, c) and genome-wide transcriptional activation potentials (Fig 4d). An additional truncation construct of p65 lacking the entire DNA-binding domain (p65-DDNA) was used as a control (Fig 4a). Bi-exponential fitting of normalized survival probability distributions from SMT (Fig 4b) revealed that both TAD mutants showed fractions of long-binding events (~4%) comparable to those observed with wt-p65 (Fig 2c,d). Moreover, the durations of such binding events was similar to those found for wt-p65, ~4 – 6 s (Fig 4c). However, both p65-ΔTAD1 and p65-ΔTAD scored as loss-of-function mutants (that is, z-score ~ 0 and median log_2_FC < 0; Fig 3b,c) as their overexpression in HeLa cells led to significantly lower levels of overall target gene transcription. Thus, despite comparable *t_b_*, truncation of TAD domains significantly impaired transcriptional activation (Fig 3b,c; Fig 4d). Nevertheless, transcriptional activation potentials of both p65 deletion mutants scored higher than the control construct p65-ΔDNA, indicating that a residual transcriptional activation potential was retained in TAD truncation mutants.

**Fig 4.**
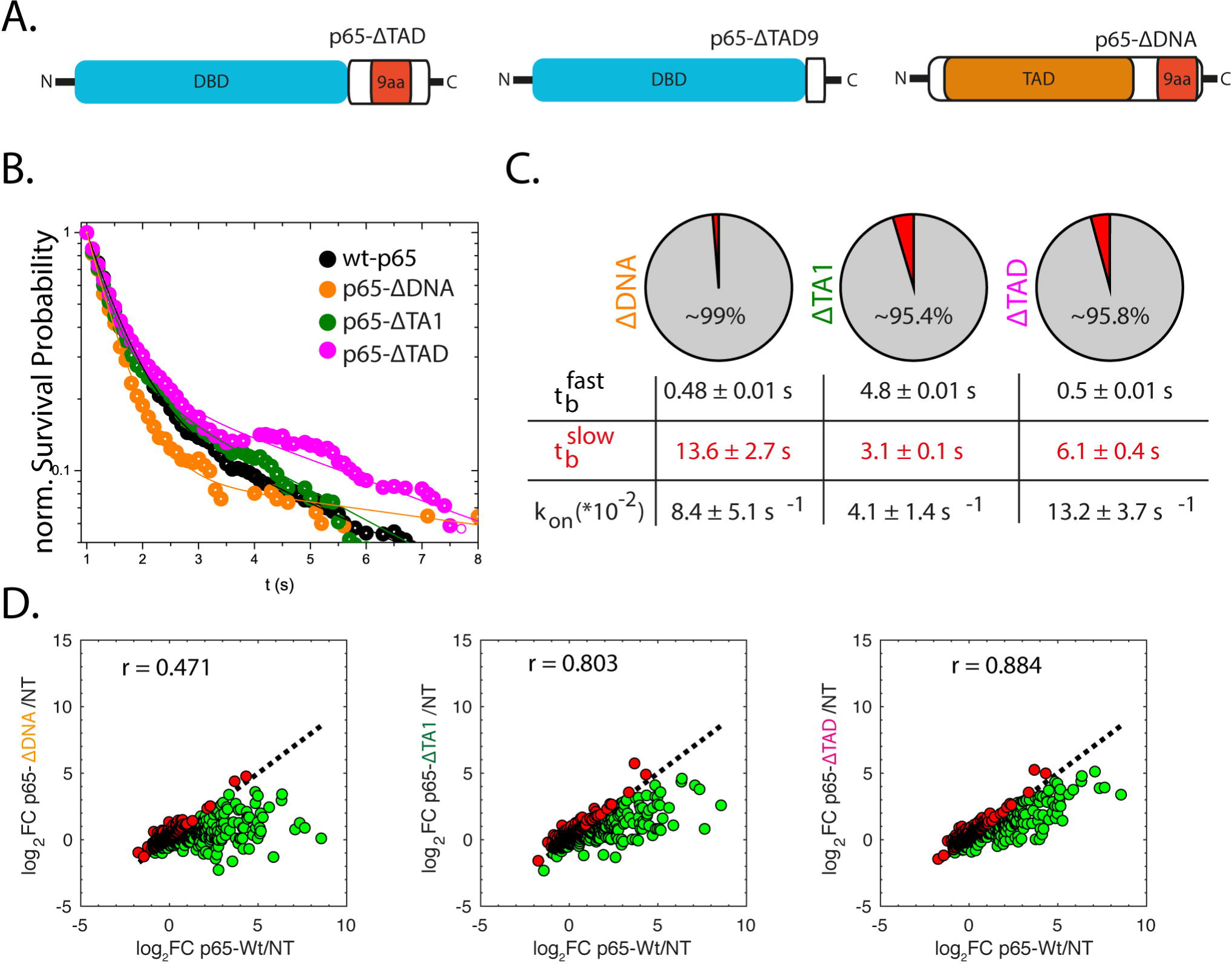
**DNA binding and transcriptional activity of p65 truncation mutants. (A)** Schematic overview of p65 truncation mutants. (**B**) Normalized survival probability (1-CDF plot) plots of the DNA-bound fraction for wt-p65 as well as the transactivation mutants ΔTA1 and ΔTAD as well as a mutant with removed DNA-binding domain (ΔDNA). The distributions were fitted using a bi-exponential function revealing the fast (t_b_fast) and slow (t_b_slow) DNA binding times. (**C**) As for the DNA affinity mutants, we found k_on_* to be in a similar range for all the tested constructs. (**D**) RNA-Seq analysis revealed very low residual transcriptional activation of p65-ΔDBD as evident by logFC ratio = −0.45. The two transactivation mutants showed good correlation with wt-p65 (r ~ 0.8) but at strongly reduced transcript abundance resulting in logFC ratio around −0.25.

The revealed relationship between transcriptional activation potential, *in vitro* p65-DNA affinity, and the duration of long-lasting DNA binding events, 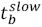, is summarized in Fig 5a. Note that the p65-DDNA mutant included in this plot was assigned an arbitrarily low relative *K_D_*~10^−5^. We observed that the median *log_2_FC* ratio of p65 DNA-binding affinity mutants correlated with 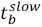. This linear dependence is recapitulated when considering correlations between the *in vitro* binding affinity (relative _9_), and 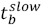. Notably, the p65-KKAA mutant appeared similar to p65-DDNA both in terms of median *K_D_* ratio and 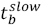.

**Fig 5.**
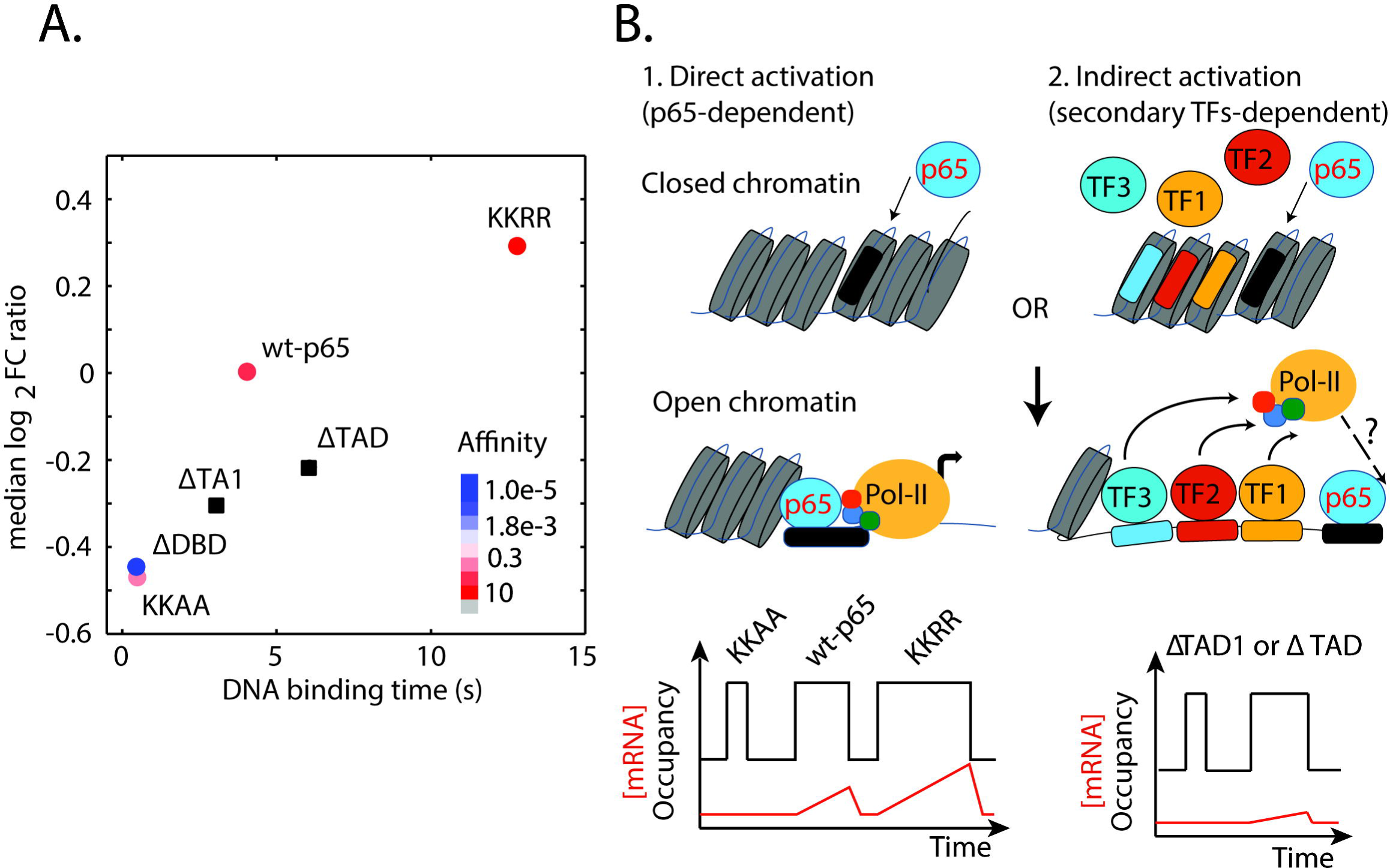
**Correlation between p65 mutants’ transcriptional activity, DNA-affinity and binding time. (A)** The median log_2_ FC ratio as retrieved from RNA-Seq data is plotted against *t*_b_*slow*. Note that DDNA affinity has been assigned to an arbitrarily low value. (**B**) Working model for p65 mediated transcriptional activation. (1) P65 can act as a pioneering TF, open the chromatin and bind its consensus DNA sequence. Transcriptional activation can then be initiated although the exact mechanism of RNA pol-recruitment remains unclear. Following this model, the DNA binding time would correlate with the transcriptional output, while removal of TADs would not affect the complex stability. (2) An important extension of this model as suggested by our data is that TADs are required to efficiently translate p65 DNA binding into transcriptional output presumably through the recruitment of protein co-factors.

Considering the impairment of protein-protein interactions induced in p65-ΔTAD1 and p65-ΔTAD mutants, we observed that both variants displayed 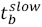 values surprisingly similar to wt-p65 (~4 *s*). Both truncations provoked significantly lower median *log_2_FC* ratios as compared to wt-p65, thus scoring much lower transcriptional activation potentials. However, it should be noted that deletion of one or both TADs did not fully abolish p65 mutants’ capability to trigger transcriptional activation, as demonstrated by comparing the median *log_2_FC* ratios of p65 truncation mutants and p65-ΔDNA, considered here as a negative control of both DNA-binding and transcriptional activation.

## Discussion

The relationship between TF DNA binding time and downstream transcriptional activation is fundamental to understanding the mechanism of gene expression and its regulation. In this work, we investigated two fundamental questions: (*i*) How does the binding time change when TF-DNA or TF-co-regulator interactions are modified or abrogatedŒ (*ii*) What are the downstream effects on gene activationŒ We found that the contribution of protein-protein interactions to p65-DNA binding stability was negligible at the genome-wide scale, since p65 truncation mutants lacking the TADs show binding times *t_b_* and bound fractions *BFs* comparable to the wild-type form of p65, (wt-p65). The model previously described for the interferon-beta (IFN-β) locus assigns a prominent function to protein-protein interactions for the stability of the protein complex – the enhanceosome – formed by p65 and associated TFs. According to this model, preventing p65 from interacting with co-regulators of the transcriptional machinery should result in shorter interactions between p65 and target chromatin binding sites. On the contrary, our results indicate that p65 *t_b_* remains largely unaffected by the absence of one or both p65-TADs. We note that heterodimers of p65 and endogenous p50 likely maintain some degree of protein-protein interactions. Nevertheless, we found that removing TADs from p65 significantly affects gene transactivation, even if it is not completely abolished. A possible interpretation of our findings assigns to TADs the general role of translating stable p65-DNA binding interactions into productive transcriptional events. Importantly, the negligible impact of TAD-mediated protein-protein interactions on p65-DNA binding stabilization, we probed at the genomic-scale, does not rule out the validity of the enhanceosome model described for the single IFN-β locus. The specific promoter architecture likely determines the stabilizing contribution of protein-protein interactions at a locus-specific scale, whereas any genomic scale-recorded readout may average these differences out.

Other studies have also challenged the enhanceosome model. One alternative proposes that protein-protein interactions between p65 and downstream components of the transcriptional machinery were proposed to actively evict p65 from chromatin, showing that the nature of the stabilizing contribution may depend on the specific components recruited to a promoter [12]. However, our findings do not recapitulate these experimental results, since wt-p65 molecules displayed similar *t_b_* as those recorded for both ΔTAD-mutants. The difference between our results and these previous measurements is again one of genetic context: we used native genes as opposed to arrays of multiple p65 binding sites stably integrated into the genome. Thus, our results more directly address the question of the effect of protein-protein interactions at the genomic scale.

In addition to the activation mechanism relying on the stabilization of protein-protein interactions, p65 is capable of triggering transcriptional activation more indirectly [9, 11]. According to this indirect model of p65-dependent gene activation, p65 can act as a “pioneer TF” that promotes chromatin opening, making adjacent regulatory elements accessible to secondary TFs. Following binding, transcriptional activation may be elicited, although the exact mechanism of RNA Pol-II recruitment at these sites has not yet been elucidated (Fig 5B). A first, important consequence of this model is that the removal of TADs is not predicted to affect the stability of p65 binding to target regulatory elements, since p65 can still undergo DNA binding through its unaltered DBD. This insight constitutes the main achievement of the present work, as demonstrated above. A second, more subtle implication of the model concerns the detectable levels of transcription when p65 lacks TADs. As previously shown [21], truncation mutants can still trigger transcriptional activation at specific loci. Notably, this is consistent with our results, since we detected residual gene expression levels when either p65-ΔTAD1 or p65-ΔTAD1/2 were overexpressed in our cells.

An additional intriguing finding of the present study concerns the positive correlation between p65 binding time, DNA-binding affinity and transcriptome-wide gene expression. This result recapitulates predictions of the “clutch model” [10], extrapolated from biochemical evidence. According to this model, longer TF binding times should yield higher expression levels of target genes. Interestingly, this model was recently challenged by two key studies, but found to hold for both the transcriptional activator p53 [11] and artificial repressor-like effectors [9]. Although the present study was carried out by overexpressing recombinant p65 constructs in HeLa cells, our experimental results recapitulate the expected trend of transcriptional levels [19, 21]. Future studies may consider genome editing approaches to avoid overexpression and remove contributions from endogenous p65.

A fundamental aspect of the present study is that we combined SMT with RNAseq to inspect how p65 binding kinetics correlate with the regulation of gene expression at a genomic scale. Our approach sheds new light on the mechanistic role of p65 trans-activation domains in regulating p65-DNA binding kinetics and the relative transcriptional outcome. We gained also new insights on how p65-DNA binding affinity may tune gene expression, underpinning an emerging model of transcriptional regulation in higher eukaryotes.

## Materials and Methods

### Cells and Plasmids

Human HeLa cells were cultured in full-supplemented DMEM (high-glucose DMEM, Gibco; 10% vol/vol fetal bovine serum, FBS, Gibco; 1% vol/vol of penicillin/streptomycin mix, Gibco and 1 mM L-glutamine, Gibco). For regular HeLa subculturing, a subcultivation volumetric ratio of 1:5 – 1:7 was used every 24-48 hours, respectively. HeLa cells were transiently transfected with Lipofectamine 3000 (ThermoFischer Scientific). Cells were seeded in 6-well plates at a density of ~2.0*10^5^ cells/well about 16-20 hours before transfection was performed in antibiotic-free, full-supplemented DMEM. 7.5 μL of Lipofectamine 3000 and 5.0 μg of plasmid DNA were then diluted each in 125 μL of room-temperature OptiMEM (Gibco) in two distinct 1.5 mL Eppendorf tubes. Diluted DNA was supplemented with 10 μL (2 μL/μg of DNA) of P3000™ reagent, mixed, and added to diluted Lipofectamine 3000 reagent. Complexes were incubated 15 minutes at RT and evenly distributed on 2.0 mL of fresh, full-supplemented DMEM medium without antibiotics. For microscopic imaging, 10-12 hours later, cells were labelled by adding 0.1-0.5 nM JF549 (L. Lavis, Janelia) in phenol-red free DMEM (LifeTechnologies) supplemented with 10 % vol/vol FBS for 30 minutes at 37°C/5% CO2. HeLa cells were then washed 3 times for 20 minutes with phenol-red free complete DMEM to remove excess fluorophore. The mammalian expression vector encoding the Halo- and FLAG-tagged, wild-type human p65 (pCI-neo-p65-Halo-FLAG) was originally obtained from Promega and described in [13]. Point mutants (KKAA, KKRR) and deletion mutants (ΔDNA, ΔTAD and ΔTA1) were generated by mutagenesis directly from pCI-neo-p65-Halo-FLAG. The QuickChange Site-Directed Mutagenesis kit (Stratagene) was used to make point mutations within the wild-type p65 coding sequence and generate KKAA, KKRR. To generate ΔDNA and ΔTAD deletion mutants, an overlap extension PCR protocol was used.

### Quantitative real-time PCR (qRT-PCR)

qRT-PCR was used to functionally validate the p65 construct encoded in the pCI-neo-p65-Halo-FLAG expression vector. HeLa cells were seeded in 6-well plates and either transfected or not with pCI-neo-p65-Halo-FLAG plasmid using Lipofectamine 3000. 24 hours later, cells were quickly rinsed with pre-warmed, sterile PBS before performing serum-starvation for 4 hours. Cells were then either treated or not with 20 ng/mL human TNF-α (Sigma) for 30 minutes at 37°C/5% CO2. After stimulation, cells were quickly rinsed twice in ice-cold PBS and total RNA was extracted (RNAeasy Mini kit; Qiagen). Briefly, cells were directly lysed in wells using 350 μL RLT buffer supplemented with β-mercaptoethanol (β-MeOH). Lysates were combined with an equal volume of 70% vol/vol ethanol diluted in DEPC-water and loaded into provided silica mini-columns. After processing samples with 700 μL RW1 buffer and twice with 500 μL of RPE buffer, total RNA was eluted in 30 μL of nuclease-free water. Samples were stored at −80°C until use. RNA samples were retro-transcribed with the SuperScript™ II Reverse Transcriptase (SuperScript™ II RT; ThermoFischer Scientific). 250 ng of random primers (RPs; ThermoFischer Scientific) were combined with 1 μg of total extracted RNA from the previous step and 1 μL of dNTPs mix (10 mM each; ThermoFischer Scientific) in 0.2 mL sterile plastic PCR tubes. Reactions were incubated 5 minutes at 65°C and after a brief centrifugation, each sample was added with 4 μL of 5X First-Strand Buffer, 2 μL of 0.1 M Di-thio-threithol (DTT) and 1 μL of RNasin^®^ (Promega). Tubes were incubated at 25°C for 2 minutes and 1 μL (200 units) of SuperScript™ II RT was added. Tubes were allowed to incubate at 25°C for 10 minutes and then at 42°C for 50 minutes. Samples were stored at −20°C until use. Quantitative analysis of p65-target genes *NFKBIA* and *Ccl2* was performed with the support of the Gene Expression Core Facility of the EPFL. Briefly, an automatic pipetting system (Hamilton) was used to combine retro-transcribed cDNA templates with primers specific for target genes *NFKBIA* (FW: 5’-ATGTCAATGCTCAGGAGCCC-3’, RV: GACATCAGCCCCACACTTCA-3’ and *Ccl2* (GeneCopoeia) and four additional housekeeping genes (β-glucoronidase, gusB; β-actin, actB; Eukaryotic elongation factor 1-alpha, eEF-1α; and TATA-binding protein, tbp) in a 384 wells-plate. Three technical replicates were measured for each biological condition. For each qPCR reaction, 3.5 μL of forward and reverse primers premix (200 nM final concentration) were mixed with 1.5 μL of cDNA template diluted 1:5 and 5 μL of SYBR Green 2X Master Mix (Applied Biosystems^®^). 384-wells plates were briefly centrifuged and sealed before performing real-time quantitative PCR with an ABI Prism 7900 Real-time PCR machine (Applied Biosystems). To interpret the data, the threshold cycle (Ct) values obtained with the SDS software (Applied Biosystems) were imported into qBase, a Visual Basic Excel based script for the management and automated analysis of qPCR data for further analysis [22]. Ct values were transformed to normalized relative quantities (NRQs) assuming a gene-amplification efficiency of 2 (i.e. equivalent to 100%). This application for Microsoft Excel allows gene expression quantification relying on multiple reference housekeeping genes. NRQs values were reported as averages of three biological replicates ± standard-error of the mean (SEM).

### Single molecule Imaging

Single-molecule acquisitions to determine p65 binding kinetics were conducted on an Olympus IX81 inverted microscope equipped with a 100x oil-immersion objective lens (Olympus, N.A. = 1.49) and with an air-stream stage incubator (Okolab UNO, Stage Mad City Labs Z2000) that kept cell samples at 37°C and 5% CO2. The setup for single-molecule microscopy was based on an inclined illumination (HILO) scheme to reduce the background signal originated from out-of-focus molecules [17] and arranged as previously described [18]. Specimen was mounted on a piezoelectric stage enabling selection of the focal plane without modifying the position of the objective. Such a configuration allowed us to adjust the focal plane so that to lie approximately in a middle section of the cell nucleus. Single-molecule stacks (300 frames/stack; 128 × 128 pixels; 18.56 × 18.56 μm^2^) were acquired by strobing the excitation 561 nm laser (Qioptiq iFlex Mustang). Specifically, to record the binding time (*t_b_*) of p65 variants, the EM-CCD camera (Evolve 512; Photometrics) and the laser were synchronized by means of a pulse generator in order to avoid photobleaching when the camera shutter was closed, using an integration time (*t_int_*) of 5 ms and a gap time (*t_gap_*) of 95 ms (referred in the following as “slow movies”). Single-step displacement analysis (ssd; see next section) and bound-fraction (*BF*) were computed out of “fast movies”, where *t_int_* = 5 *ms* and *t_gap_* = 15 *ms*. An irradiation intensity of ~1 kW/cm2 was used for both settings. Image stacks were collected using μManager open source microscopy software, setting the EM-CCD camera electronic multiplier (EM) gain to 300 AU.

### Image Analysis

Movies collected for each p65 construct were analyzed using a Matlab routine (MatlabTrack_v5.03) described in [18]. Individual frames were processed with a band-pass filter using a lower threshold of 1 pixel (equivalent to 145 nm) and a higher threshold of 5 pixels both to smooth the diffraction limited spots corresponding to single molecules and suppress pixel noise. Localization of fluorescent peaks was carried out by using a dedicated algorithm implemented within MatlabTrack_v5.03 using an intensity threshold of 500-700 AU, visually adjusted according to the noise level of the movie. These threshold values allowed us to discard dim peak intensities putatively corresponding to out-of-focus molecules. Tracking was performed by using MatlabTrack_v5.03 which implemented the Matlab version of the Crocker and Grier algorithm [23].

We analyzed “slow movies” (*t_gap_* = 95 ms) to estimate the distribution of p65 residence times: to this scope we allowed for a maximum displacement between consecutive frames of 5 pixels (725 nm) to selectively identify slowly moving or immobile molecules. To account for blinking of the fluorophore we allowed an arbitrary gap-length of 3 frames. We discarded tracks shorter than 2 frames. A more stringent selection of putative p65 binding events was performed by using an additional filter implemented in MatlabTrack_v5.03. Specifically, we retained only molecules displacing shorter than 3 pixels (435 nm) and longer than 10 frames (1 s) as previously described [18] by comparison with immobile H2B molecules (S2 Fig). This allows to discard slowly mobile p65 molecules that might otherwise be erroneously interpreted as bound.

Trajectory were calculated out of individual movies collected for each p65 mutant (10-12 movies per condition) using MatlabTrack_v5.03. The duration of each track was assumed to be equal to the time the molecule stays bound while unbleached, i.e. the binding time, *tb*=1/*koff*, where *koff* corresponds to the kinetic dissociation rate of each detected single molecule. Each distribution corresponding to the different p65 variants was normalized against the trajectory length distribution of H2B to account for photobleaching.

In order to resolve the fast and slow kinetic components *kfast* and *kslow*, the 1-cumulative distribution function (1-CDF) histogram for different p65 variants was calculated based on the binding time of each individual tracks. Values corresponding to calculated 1-CDF distributions were then exported in OriginPro9 (OriginLab) and fitted according to either a mono- or bi-exponential decay function. Goodness of fitting was evaluated by the chi2 and selection of the fitting function was based on a nested-model *F*-test as implemented in OriginPro9 (for details: www.originlab.com/doc/Origin-help/PostFit-CompareFitFunc-Dialog).

The bound fraction (BF) was calculated by performing the single-step displacement (ssd) analysis as described in [18] using “fast movies” collected with *t_gap_* = 15 ms. Briefly, the probability density distribution (*r*) of displacing a distance between *r* and *r*+Δ*r* in the time Δ*t* between two consecutive frames in our single-molecule movies of Halo-tagged p65 was fit by a *n*-component diffusion model:

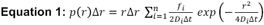

where *Di* are the diffusion coefficients for each of the species and *fi* are the fractions of molecules with diffusion coefficient *Di*, with 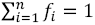. We found that a three-component diffusion model provided adequate fitting of the experimental data and the slowest diffusion component *D*1 matched the average diffusion coefficient measured for the chromatin-bound histone subunit H2B (~0.04 μm2s-1). Therefore, in the following we will report *f*1 as equivalent to the BF for both wt- and mutant p65.

### Transcriptome-wide RNA-Sequencing

The RNAseq experiment was run through the genomic technologies facility of the University of Lausanne and the bioinformatics and biostatistics core facility of the EPFL. Purity-filtered reads were adapted and quality trimmed with Cutadapt (v. 1.3, [24]) and filtered for low complexity with seq_crumbs (v. 0.1.8). Reads were aligned against *Homo sapiens v.* GRCh38 genome using STAR (v. 2.4.2a, [25]). The number of read counts per gene locus was summarized with htseq-count (v. 0.6.1, [26]) using *H. sapiens v.* GRCh38 Ensembl 82 gene annotation. Quality of the RNA-seq data alignment was assessed using RSeQC (v. 2.3.7, [27]). Reads were also aligned to the *H. sapiens v.* GRCh38 Ensembl 82 transcriptome using STAR (v. 2.4.2a, [25]) and the estimation of the isoforms abundance was computed using RSEM (v. 1.2.19, [28]). Statistical analysis was performed for protein-coding genes and long non-coding genes genes in R (R version 3.1.2). Genes with low counts were filtered out according to the rule of 1 count per million (cpm) in at least 1 sample. Library sizes were scaled using TMM normalization (EdgeR v 3.8.5; [29]) and log-transformed with *limma voom* function (R version 3.22.4; [30]). Differential expression was computed with *limma* [31] by fitting data into a linear model, extracting the contrasts for all pairwise comparisons of transfected vs Non-transfected (NT). A moderated F-test was applied and the adjusted p-value computed by the Benjamini-Hochberg method, controlling for false discovery rate (FDR). Genes displaying a FDR < 0.1 were selected as differentially expressed. Data analysis performed as described above identified 1080 differentially expressed genes at FDR 10%.

However, the transcriptional activity of the different p65-Halo variants was assessed using only a subset of 215 differentially expressed genes selected as direct binding targets of p65 on the basis of the ENCODE ChIP-seq deposited information (Dr. Jacques Rougemont; Bioinformatics and biostatistics core facility, EPFL). To calculate the intersection between differentially expressed genes based on RNAseq analysis and the ENCODE ChIP-seq database, we first identified find the differentially expressed gene (RNASeq) coordinates using *genrep4humans.py* assembly hg19. Then we defined regions of interest on the forward strand from Gene_Start-2000 to Gene_End and on the reverse strand from Gene_Start to Gene_End+2000. We use *bedtools intersect* to get the intersection of the peaks and the regions promoter+gene. We find at least one peak in 19.9% of the differentially expressed genes: 215 / 1080.

We used *venn_mpl.py* from *pybedtools* to plot a Venn diagram of genomic regions. We plotted the intersection between the promoter + gene region and promoter regions of the differentially expressed genes and the ChIPSeq Peak regions using the hg19 assembly. We performed the NFKB1_REL_RELA.P2 motif search (obtained from the Swissregulon Database at http://swissregulon.unibas.ch/fcgi/wmŒwm=NFKB1_REL_RELA.p2&org=hg19) in the promoter regions of the selected genes (on the forward strand Gene_Start+/-2000 and on the reverse strand Gene_End+/-2000). In order to retrieve the FASTA sequence, we used *bbcfutils*, a collection of tools used at the Bioinformatics & Biostatistics Core Facility, EPFL, Lausanne, Switzerland. More precisely, the *genrep4humans.py* script gives the gene coordinates with the assembly hg38, (https://github.com/bbcf/bbcfutils/blob/master/Python/genrep4humans.py). Average expression levels of genes scoring at least one ChIP-seq peak in either the promoter or coding sequence were visually represented in a single heat map calculated in MatLab. The heat map displays the expression levels of the 215 genes identified from the ENCODE ChIP-seq database obtained for each p65 mutant and corrected for the NT sample. For each p65 mutant, we plotted the relative expression (log2 fold-change; log2FC) of the wild-type p65-transfected condition on x-axis and mutant p65-transfected condition on y-axis, both compared to the wild-type non-transfected condition (NT). We computed the correlation coefficient (r) and the median log2 FC over all genes. The later could then be used to classify p65 mutants as either loss- or gain-of function.

### Fluorescence recovery after photobleaching (FRAP)

At 12-16 hours post-transfection, Hela cells were labeled with 5 μM OregonGreen Halo Ligand (Promega) for 30 minutes at 37°C/5% CO_2_. Cells were then extensively washed in pre-warmed phenol red-free DMEM so that to be sure to have eliminated the majority of unbound fluorescent ligand. FRAP experiments have been carried out with the Leica SP8 confocal fluorescence microscope using an oil-immersion PLAN-APOCHROMAT 60X objective. Fluorescence was excited with an Argon laser set at 80% of its total power. Pre- and post-bleach images (256×6 pixels) were acquired with a pinhole aperture set to 2 Airy-units using bidirectional scanning mode for faster acquisition. A total of 50 pre-bleach and 500 post-bleach frames were collected at 0.2% of AOTF and a zoom factor of 8 that resulted in a final pixel size of 180 nm. The delay time between successive frames was 69 ms. Bleaching was obtained using the 488 nm Argon laser set at maximum power and the zoom-in option implemented in the TCS SP8 FRAP module (one bleaching frame only).

Regions of interest (ROIs) corresponding to the photobleached area, the whole nucleus and the background region were manually segmented in Fiji (https://imagej.nih.gov/ij/) for each recorded cell. FRAP curves were then calculated and normalized using FRAPAnalyser 2.0 (http://actinsim.uni.lu/). Double exponential fitting was performed according to:

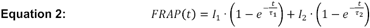

which was implemented in the FRAPAnalyser 2.0 τ_1_ and τ_2_ are the time-constants corresponding to the fast and the slow component, respectively, calculated as τ = 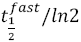 and 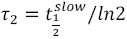, being 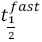 and 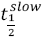 the half-times of recovery of the fast and slow fractions.

The global half-time of recovery 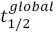 corresponds to the 50% of fluorescence signal recovery. The value of *FRAP* 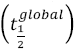 can be computed as:

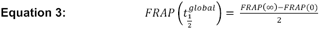

Given that: 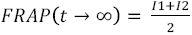 and 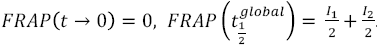. Therefore, when 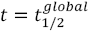 we have that:

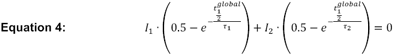

To compute 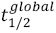, the equation reported above is solved numerically in R using the function *uniroot* (see S1 Table).

### Extraction of ectopically expressed p65-Halo-FLAG from HEK293 cells

Suspension-adapted HEK293 cells were routinely maintained in serum-free ExCell 293 medium (SAFC Biosciences, St. Louis, MO) with 4 mM glutamine as described [32] in a shaking ISF-4-W incubator (Kühner AG, Birsfelden, Switzerland) at 37°C in the presence of 5% CO_2_ at the Protein Expression Core Facility of the EPFL (In collaboration with Dr. D. Hacker). HEK293 cells were transfected with pCI-neo-p65-Halo-FLAG as described in 24 hours post-transfected cells (~10^9^) were harvested by centrifugation (500×g, 5 minutes at RT) in two 50-mL Falcon tubes. Supernatant was filtered and maintained in the cell incubator at 37°C with 5% CO_2_ to perform TNF-a stimulation. Cell pellets were pooled together in a single 50-mL Falcon tube and resuspended in 40 mL of the original pre-equilibrated cell culture medium supplemented with 20 ng/mL of human TNF-a (Sigma). The cell suspension was incubated for 30 minutes at 37°C in an orbital shaker (180 rpm). Stimulated HEK293 cells were then pelleted (2000×g for 5 minutes at 4°C) and resuspended in ice-cold PBS added with phosphatase (Sigma) and protease inhibitors (COMPLETE™; Roche). Cell washing with supplemented-PBS was repeated once and the cell pellet was frozen at −80°C for 1 hour to help protein releasing from cells due to a facilitated plasma-membrane rupture out of a freeze-thaw cycle. Thawed cells were added with ~5 packed cell volume (pcv) of PBS with 1% vol/vol phosphatase inhibitors (Sigma) and 1 mM DTT and pelleted at 1’800×g for 5 minutes at 4°C. Cell pellet was resuspended in ~3 pcv of hypotonic buffer (10 mM HEPES, pH 7.9 at 4°C; 1.5 mM MgCl_2_; 10 mM KCl; immediately before use add protease inhibitors (1 COMPLETE™ table/10 mL of buffer) and 1 mM DTT) and incubated on ice for 10 minutes. Cells were then homogenized with a glass, ice-cold 15-ml Dounce homogenizer (pestle B, 28 strokes on ice; Kimble Chase). This step disrupts the majority of cells membranes but keeps nuclei intact. Nuclei were then pelleted (3’300×g for 15 minutes at 4°C) and resuspended in 1 packed-nuclear volume (pnv) of low-salt buffer (20 mM HEPES, pH 7.9 at 4°C; 25% glycerol; 1.5 mM MgCl_2_; 0.2 mM EDTA; supplement with protease inhibitors and 1 mM DTT prior to use). Nuclei were then dispersed thoroughly with the 15-mL Dounce homogenizer (pestle B) while adding 5 M NaCl dropwise up to a final concentration of 420 mM to allow chromatin-bound proteins to be extracted from nuclei. Nuclear lysates were incubated for 30 minutes at 4°C on a rotating wheel and ultracentrifuged (100’000×g for 1 hour at 4°C). The supernatant was then collected and diluted with one volume of hypotonic buffer supplemented with 1 mM DTT, 20% vol/vol glycerol, 0.2% NP-40 alternative and phosphatase/protease inhibitors.

### Pull-down of extracted p65-Halo-FLAG

The recombinant p65-Halo-FLAG protein was purified from nuclear crude extracts by performing a pull-down with anti-FLAG M2 magnetic beads (Sigma). 2.0 mL of beads were washed three times in 5 mL of equilibration buffer (10 mM HEPES, pH 7.9 at 4°C; 10 mM KCl; 1.5 mM MgCl2; 200 mM NaCl; 0.1% vol/vol NP-40 alternative; 10% vol/vol glycerol) and collected through the magnet. The nuclear crude extract (~6 mL) was then added to beads together with 14 μM of JF549 fluorescent Halo-ligand (from Dr. L. Lavis) and incubated ON at 4°C on a rotating wheel. After extensive washing in elution buffer (10 mM HEPES, pH 7.9 at 4°C; 200 mM NaCl; 0.1% vol/vol NP-40 alternative; 1 mM DTT; 1 mM EDTA; 10% vol/vol glycerol freshly supplemented with protease and phosphatase inhibitors) to remove unbound proteins and excess fluorophore, beads were incubated in elution buffer supplemented with 100 μg/mL of FLAG peptide (Sigma) for 1 hour at 4°C on a rotating wheel. The supernatant was then collected and stored at 4°C until use. The elution step was performed three times and supernatants were pooled together and concentrated in Centricon 10 kDa MWCO centrifuge filters at 5000 × g for ~2 hours at 4°C.

### SDS-PAGE and Western Blot

The eluted p65-Halo-FLAG protein concentration was determined by performing denaturing sodium dodecyl sulphate-polyacrylamide gel electrophoresis (SDS-PAGE) against known amounts of bovine serum albumin (BSA) standards, followed by Coomassie staining (SimplyBlue™ SafeStain; Thermo Fischer Scientific). Variable volumes of eluted p65-Halo-FLAG (0.5 μL, 5 μL and 10 μL) and BSA standards (0.2 μg, 0.5 μg, 1.0 μg, 1.5 μg, 3.0 μg and 4.0 μg) were denatured in 1x Laemmli Sample Buffer (Alfa Aesar) and boiled for 5 minutes at 95°C. Samples (20 μL final volume) were separated using 12% SDS-PAGE prepared from stock 37.5:1 polyacrylamide:bis-acrylamide solution (Fischer Scientific) and run at 120 Volts for ~1 hour in Tris-Glycine running buffer (25 mM TrisCl; 250 mM glycine; 0.1% SDS) using a MiniProtean™ System (Biorad). For Coomassie staining, the minigel was rinsed three times with ~100 mL deionized water and ~20 mL of blue stain were added and incubated ON. The minigel was destained 2 hours with 100 mL of water. The final protein concentration was ~47 ng/μL (~470 nM) as estimated from densitometry (ImageJ).

For Western Blot, samples preparation and electrophoresis were performed as described above. Proteins were ON-transferred to a nitrocellulose membrane (Protran™ Hybond ECL; GE Healthcare) at 4°C in Towbin transfer buffer (25 mM TrisCl; 192 mM glycine, pH 8.3; 20% methanol and 0.1% SDS) at 100 Volts using the MiniProtean™ transfer cassette (Biorad). Membranes were then blocked in non-fat dry milk (5% w/vol; Biorad) for 1 hour at RT and probed with mouse monoclonal IgG_1_ anti-p50 antibody (1:200; Santa Cruz) ON at 4°C in TBST (20 mM TrisCl pH 7.5; 150 mM NaCl; 0.1% Tween.20) supplemented with 5% w/vol non-fat dry milk. Filters were then washed 3 times for 15 minutes each with TBST and probed with a sheep anti-mouse, peroxidase-labelled antibody (Amersham) for 45 minutes at RT in TBST supplemented with 5% w/vol non-fat dry milk. Membranes were washed 3 times for 15 minutes with TBST and developed with ECL Plus system (Thermo Scientific). Chemiluminescence detection was carried out with a gel fluorescence scanner (ChemiDoc; Biorad). Notably, p65-Halo-FLAG was detected directly through the fluorescence emitted from the covalently-bound JF549 and, therefore, did not need to be probed with a specific antibody.

### Electrophoretic mobility shift assay (EMSA)

Synthetic HPLC-purified sense and anti-sense oligo probes encoding the consensus binding sequence of p65 were purchased from Microsynth (Microsynth AG, Switzerland; Sense-p65_κB: 5’-AGTTGAG<u>GGGGACTTTCC</u>CAGGC-3’; Anti-sense-p65_κB: 5’-GCCTG<u>GGAAAGTCCC</u>CTCAACT-3’). An Atto647N dye was attached to the 5’ of the sense-strand to visualize DNA by fluorescence detection. To make double-stranded DNA probes, a pair of sense and anti-sense oligos were mixed at 50 μM each in annealing buffer (10 mM TrisCl, pH 8.0; 1 mM EDTA and 50 mM NaCl), then annealed in a PCR machine with the following program: 95°C (3 minutes), 0.1°C/second drop to 55°C, 55°C (60 minutes), 0.1°C/second drop to 25°C as previously reported [34]. In EMSA, 0.5 μM fluorescent double-stranded DNA probe (dsDNA) was mixed with 0.1, 0.3 and 0.6 μg of purified, JF549-labeled p65-Halo-FLAG in 20 μL binding buffer (25 mM HEPES, pH 7.6; 0.1 mM EDTA; 12.5 mM MgCl_2_; 100 mM KCl; 0.01% NP-40 alternative and 10% glycerol) and incubated at 4°C for 60 minutes and then at RT for 10 minutes before loading into 1% w/vol agarose (Sigma) prepared in 1x TBE buffer (45 mM Tris-borate; 1 mM EDTA) and prerun at 120 Volts, 4°C for 30 minutes. Samples were run by electrophoresis at 150 Volts for ~1 hour at 4°C in 0.5x TBE buffer. Fluorescence signals were scanned with a Chemidoc imaging system (Biorad).

## Acknowledgements

We thank the Genomic Technologies Facility of the University of Lausanne and the Bioinformatics and Biostatistics Core Facility of the EPFL for conducting the RNA-Seq experiments and help with data analysis. The FRAP experiments were carried out in ALEMBIC, an advanced microscopy facility established by the San Raffaele Scientific Institute. This work was supported by funds from the Swiss National Science Foundation (A.C., S.M. (CR33I2_149850) D.M.S. (PP00P3_144828)), the Swiss National Center for Competence in Research (NCCR) Chemical Biology (S.M., C.S., B.F.), and an EMBO Short-Term Fellowship (A.C.).

## Author contributions

A.C., A.B. and S.M. conceived the project. All authors contributed to the design of the project. D.S., B.F., D.M. and S.M. supervised the project. A.C, C.S. and D.M. performed all experiments and data analysis. All authors contributed to writing and revising the final manuscript.

---

**S1 Fig.** qPCR of Hela cells either transfected (+p65-HT) or non-transfected (-p65-HT) with p65-Halo wild-type construct (wt-p65) in the presence (+) or absence (−) of TNF-α. Averages of normalized relative quantities (RQs) of biological triplicates ± standard deviation (SD) are shown. *\* p* < 0.05.

**S2 Fig.** (**A**) Western blot of p65-Halo purified fraction. Merged signals out of anti-p65 and anti-p50 are shown. (**B**) Electrophoretic mobility shift assay (EMSA) of wild-type p65-Halo-tagged construct (p65-HT) and its consensus oligo (ds-κB-oligo). Increasing quantities (in μg) of JF549-labeled p65-HT (green channel) and a constant amount of ds-κB-oligo (red channel) are incubated together and electrophoretically separated under native conditions. The merge of both channels shows overlapping p65-HT fluorescence with consensus oligo signal.

**S3 Fig.** (**A**) Binding events of individual p65 molecules are detected based on both spatial (435 nm) and temporal (1 s) thresholds experimentally established from imaging of immobile H2B molecules. (**B**) Analysis of bound segments of H2B (left) and wt-p65 (right), using the spatiotemporal criteria explained in panel A, assigns ~99% of H2B molecules and ~40% of wt-p65 to the ‘bound’ state.

**S4 Fig.** Normalized survival probability distributions of wt-p65 and its DNA-binding affinity mutants KKAA and KKRR. Mono- (dot line) and bi-exponential (continuous line) fitting models are compared for each binding time distribution.

**S5 Fig.** (**A**) Fluorescence recovery after photobleaching (FRAP) of wt-p65 and its DNA-binding affinity mutants. Pre- and post-bleaching snapshots of a representative nucleus overexpressing the H2B-Halo construct (top). The actual size of the bleached region is highlighted with a white rectangle. Note that H2B-Halo fluorescence does not recover, confirming that H2B-Halo is immobile in living Hela cells. Different regions of interest (ROIs) used to calculate the FRAP recovery curves are indicated with numbers (1, 2 and 3; low). **1**: bleaching ROI; **2**: reference ROI encompassing the whole nuclear area used to normalize against the actual expression levels and photobleaching; **3**: background. Representative time-points of wt-p65 fluorescence recovery are shown. Scale bar: 5 μm. Averaged normalized FRAP curves of wt-p65, DNA-binding affinity mutants and DDNA (control) collected from Hela cells stimulated as described in Methods (left). Curves obtained from double-exponential model fitting of experimental data-points (see Methods) are superimposed to estimate 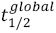. Distributions of 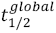 are represented as box-plots (right). The average (black square) and the median values (horizontal line) of each distribution are displayed for each box-plot together with the number (n) of measured cell nuclei. Whiskers span over the 25%-75% percentile range. (**B**) Analysis of p65 bound fraction (BF). Histogram of the BF values of wt-p65 and its mutants compares each construct to H2B bound-fraction. Single-step displacement distribution analysis (inset) and its fitting with a three-component diffusion model (black curve) to retrieve BF. The diffusion constant calculated for the DNA-bound H2B histone subunit (red dots and line) matched the slowest diffusing component of the p65 construct whose amplitude corresponds to the computed BF.

**S1 Table.** Global FRAP recovery half-times (50 % recovery) for the indicated conditions computed using Equation 3.

## References

Medzhitov R, Horng T. Transcriptional control of the inflammatory response. Nat Rev Immunol. 2009;9: 692–703. doi:10.1038/nri2634

Baeuerle PA, Baltimore D. IκB: A specific inhibitor of the NF-κB transcription factor. Science. 1988;242: 540–546.

Baeuerle PA, Henkel T. Function and activation of NF-κB in the immune system. Annu Rev Immunol. 1994;12: 141–179.

Müller CW, Van D, Sodeoka M, Verdine GL, Harrison SC. Structure of the NF-κB p50 homodimer bound to dna. Nature. 1995;373: 311–317. doi:10.1038/373311a0

Ghosh G, Van D, Ghosh S, Sigler PB. Structure of NF-κB p50 homodimer bound to a κb site. Nature. 1995;373: 303–310. doi:10.1038/373303a0

Chen FE, Huang D-B, Chen Y-Q, Ghosh G. Crystal structure of p50/p65 heterodimer of transcription factor NF-κB bound to DNA. Nature. 1998;391: 410–412. doi:10.1038/34956

Schmitz ML, Baeuerle PA. The p65 subunit is responsible for the strong transcription activating potential of NF-κB. EMBO J. 1991;10: 3805–3817.

Bhatt D, Ghosh S. Regulation of the NF-κB-mediated transcription of inflammatory genes. Front Immunol. 2014;5. doi:10.3389/fimmu.2014.00071

Clauß K, Popp AP, Schulze L, Hettich J, Reisser M, Escoter T, et al. DNA residence time is a regulatory factor of transcription repression. Nucleic Acids Res. 2017;45: 11121–11130. doi:10.1093/nar/gkx728

Lickwar CR, Mueller F, Hanlon SE, McNally JG, Lieb JD. Genome-wide protein-DNA binding dynamics suggest a molecular clutch for transcription factor function. Nature. 2012;484: 251–255. doi:10.1038/nature10985

Loffreda A, Jacchetti E, Antunes S, Rainone P, Daniele T, Morisaki T, et al. Live-cell p53 single-molecule binding is modulated by C-terminal acetylation and correlates with transcriptional activity. Nat Commun. 2017;8. doi:10.1038/s41467-017-00398-7

Bosisio D, Marazzi I, Agresti A, Shimizu N, Bianchi ME, Natoli G. A hyper-dynamic equilibrium between promoter-bound and nucleoplasmic dimers controls NF-κB-dependent gene activity. EMBO J. 2006;25: 798–810. doi:10.1038/sj.emboj.7600977

Los GV, Encell LP, McDougall MG, Hartzell DD, Karassina N, Zimprich C, et al. HaloTag: A novel protein labeling technology for cell imaging and protein analysis. ACS Chem Biol. 2008;3: 373–382. doi:10.1021/cb800025k

Grimm JB, English BP, Chen J, Slaughter JP, Zhang Z, Revyakin A, et al. A general method to improve fluorophores for live-cell and single-molecule microscopy. Nat Methods. 2015;12: 244–250. doi:10.1038/nmeth.3256

Buxadé M, Lunazzi G, Minguillón J, Iborra S, Berga-Bolaños R, del V, et al. Gene expression induced by Toll-like receptors in macrophages requires the transcription factor NFAT5. J Exp Med. 2012;209: 379–393. doi:10.1084/jem.20111569

Tay S, Hughey JJ, Lee TK, Lipniacki T, Quake SR, Covert MW. Single-cell NF-B dynamics reveal digital activation and analogue information processing. Nature. 2010;466: 267–271. doi:10.1038/nature09145

Tokunaga M, Imamoto N, Sakata-Sogawa K. Highly inclined thin illumination enables clear single-molecule imaging in cells. Nat Methods. 2008;5: 159–161. doi:10.1038/nmeth1171

Mazza D, Abernathy A, Golob N, Morisaki T, McNally JG. A benchmark for chromatin binding measurements in live cells. Nucleic Acids Res. 2012;40. doi:10.1093/nar/gks701

Schaaf MJM, Willetts L, Hayes BP, Maschera B, Stylianou E, Farrow SN. The relationship between intranuclear mobility of the NF-κB subunit p65 and its DNA binding affinity. J Biol Chem. 2006;281: 22409–22420. doi:10.1074/jbc.M511086200

Speil J, Baumgart E, Siebrasse J-P, Veith R, Vinkemeier U, Kubitscheck U. Activated STAT1 transcription factors conduct distinct saltatory movements in the cell nucleus. Biophys J. 2011;101: 2592–2600. doi:10.1016/j.bpj.2011.10.006

van E, Engist B, Natoli G, Saccani S. Two modes of transcriptional activation at native promoters by NF-kappaB p65. PLoS Biol. 2009;7.

Hellemans J, Mortier G, De P, Speleman F, Vandesompele J. qBase relative quantification framework and software for management and automated analysis of real-time quantitative PCR data. Genome Biol. 2007;8.

Crocker JC, Grier DG. Methods of digital video microscopy for colloidal studies. J Colloid Interface Sci. 1996;179: 298–310. doi:10.1006/jcis.1996.0217

Martin M. Cutadapt removes adapter sequences from high-throughput sequencing reads. EMBnet.journal. 2011;17: 10–12. doi:10.14806/ej.17.1.200

Dobin A, Davis CA, Schlesinger F, Drenkow J, Zaleski C, Jha S, et al. STAR: Ultrafast universal RNA-seq aligner. Bioinformatics. 2013;29: 15–21. doi:10.1093/bioinformatics/bts635

Anders S, Pyl PT, Huber W. HTSeq-A Python framework to work with high-throughput sequencing data. Bioinformatics. 2015;31: 166–169. doi:10.1093/bioinformatics/btu638

Wang D, Westerheide SD, Hanson JL, Baldwin AS. Tumor necrosis factor α-induced phosphorylation of RelA/p65 on Ser529 is controlled by casein kinase II. J Biol Chem. 2000;275: 32592–32597.

Li B, Dewey CN. RSEM: Accurate transcript quantification from RNA-Seq data with or without a reference genome. BMC Bioinformatics. 2011;12. doi:10.1186/1471-2105-12-323

Robinson MD, McCarthy DJ, Smyth GK. edgeR: a Bioconductor package for differential expression analysis of digital gene expression data. Bioinforma Oxf Engl. 2010;26: 139–140.

Law CW, Chen Y, Shi W, Smyth GK. Voom: Precision weights unlock linear model analysis tools for RNA-seq read counts. Genome Biol. 2014;15. doi:10.1186/gb-2014-15-2-r29

Ritchie ME, Phipson B, Wu D, Hu Y, Law CW, Shi W, et al. Limma powers differential expression analyses for RNA-sequencing and microarray studies. Nucleic Acids Res. 2015;43: e47. doi:10.1093/nar/gkv007

Muller N, Girard P, Hacker DL, Jordan M, Wurm FM. Orbital shaker technology for the cultivation of mammalian cells in suspension. Biotechnol Bioeng. 2005;89: 400–406. doi:10.1002/bit.20358

Backliwal G, Hildinger M, Chenuet S, Wulhfard S, De J, Wurm FM. Rational vector design and multi-pathway modulation of HEK 293E cells yield recombinant antibody titers exceeding 1 g/l by transient transfection under serum-free conditions. Nucleic Acids Res. 2008;36. doi:10.1093/nar/gkn423

Chen J, Zhang Z, Li L, Chen B-C, Revyakin A, Hajj B, et al. Single-molecule dynamics of enhanceosome assembly in embryonic stem cells. Cell. 2014;156: 1274–1285. doi:10.1016/j.cell.2014.01.062

Brown K, Gerstberger S, Carlson L, Franzoso G, Siebenlist U. Control of IκB-α proteolysis by site-specific, signal-induced phosphorylation. Science. 1995;267: 1485–1488.

Perez-Pinera P, Kocak DD, Vockley CM, Adler AF, Kabadi AM, Polstein LR, et al. RNA-guided gene activation by CRISPR-Cas9-based transcription factors. Nat Methods. 2013;10: 973–976. doi:10.1038/nmeth.2600

